# Novel interactions within the SIR heterochromatin complex potentiate inter-subunit communication and gene repression

**DOI:** 10.1101/2024.12.23.630195

**Authors:** Jenna Kotz, E. J. Martz, Maya Nelson, Nicole Savoie, Lauren Schmitt, Jordan States, Nathan Holton, Kirk Hansen, Aaron M. Johnson

## Abstract

Organisms with smaller genomes often perform multiple functions using one multi-subunit protein complex. The *S. cerevisiae* Silent Information Regulator complex (SIRc) carries out all of the core functions of heterochromatin. SIR complexes first drive the initiation and spreading of histone deacetylation in an iterative manner. Subsequently, the same complexes are incorporated stably with nucleosomes, driving compaction and repression of the underlying chromatin domain. These two distinct functions of SIRc have each been characterized in much detail, but the mechanism by which the dynamic spreading state switches to stable compaction is not well-understood. This incomplete knowledge of potential intra-complex communication is partly due to a lack of structural information of the complex as a whole; only structures of fragments have been determined to date. Using cross-linking mass spectrometry in solution, we identified a novel inter-subunit interaction that physically connects the two states of SIRc. The Sir2 deacetylase makes direct interactions with the scaffolding subunit Sir4 through its coiled-coil domain, which also interacts with the Sir3 compaction/repression subunit. Within the hub of interactions are conserved residues in Sir2 that can sense deacetylation state, as well as amino acids that likely diverged and co-evolved to interact with Sir4, promoting species-specific functions. Mutation of this interaction hub disrupts heterochromatic repression, potentially by disrupting a conserved mechanism that communicates completion of deacetylation to switch to compaction. Our work highlights how a single multi-functional chromatin regulatory complex can stage a step-wise mechanism that requires a major transition in activities to achieve epigenetic gene repression.

## INTRODUCTION

Budding yeast heterochromatin is a powerful simplified model to study chromatin-mediated gene silencing[1, 2]. The SIR complex is a stripped-down heterochromatin machine, necessary and sufficient for transcriptional repression. Yeast heterochromatin shares a basic mechanism of formation with heterochromatin in all eukaryotes separated by a dynamic phase: recruitment of a histone modifying complex to a region targeted for silencing, nucleation of heterochromatin by histone modification and sequence-independent spreading of silencing across larger regions of the genome, dependent on binding of a heterochromatin protein to the modified histone; and a stable phase characterized by compaction of the genome (Figure 1A). The SIR complex incorporates Sir2, an NAD^+^-dependent histone deacetylase (HDAC), conserved in all eukaryotes, whose activity is required for silencing[3, 4]. The SIR complex (Fig. 1B,C) is scaffolded by the Sir4 subunit, and Sir3 reads the histone modification pattern and allows the SIR complex to compact nucleosomes only when they have certain modifications erased, including de-acetylation of histone H4 lysine 16 (H4K16), which is critical for gene silencing. SIR heterochromatin stably and heritably silences the genes that dictate mating-type cell differentiation and genes in subtelomeric regions. A wealth of genetic and biochemical data from the SIR system has developed one of the richest models of heterochromatin regulation that is ripe for a new level of mechanistic structure-function analysis. The in vitro SIR heterochromatin biochemical system that we have developed in budding yeast[5] has established a model to study a stripped-down silencing mechanism with unprecedented detail.

**Figure 1.**
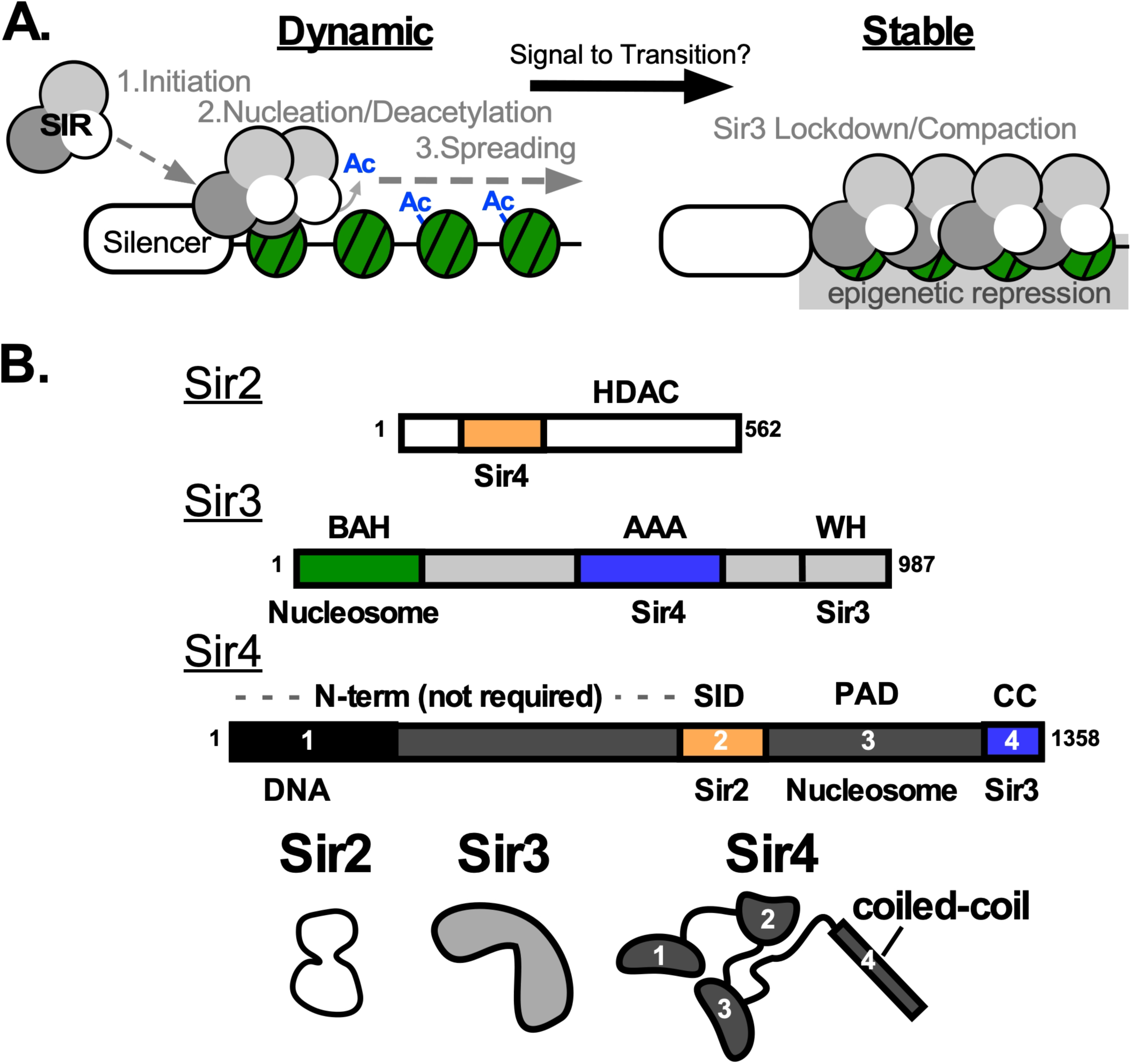
Mechanism and architecture of the SIR system. A) SIR heterochromatin works in two primary modes: 1) a dynamic mode of recruitment of the SIR complex to target sites on chromatin which then leads to nucleation of the histones by deacetylation and spreading of this part of the mechanism to contiguously deacetylate the region, 2) a stable mode where the Sir3 subunit coordinates compaction of chromatin by the SIR complex and stable incorporation of the complex into the heterochromatin domain. After initiation, the mechanism spreads and represses in a sequence-independent epigenetic manner, where the extent of spread of the domain can be inherited. As noted, a major question that remains in the field is how the singular, muti-functional SIR complex switches from one mode to the next, since the Sir2 active site directly competes with the Sir3 lockdown site. B) Domain structure of the core SIR subunits. Sir2, an NAD+-dependent histone deacetylase (HDAC), has a core catalytic HDAC domain downstream of a region that interacts with the Sir4 subunit. Sir3 has an N-terminal bromo-adjacent homology (BAH) domain that binds to a nucleosome that is deacetylated at histone H4 K16. A non-canonical AAA ATPase domain, missing catalytic residues, is in the middle of the protein and this region binds Sir4. At its C-terminus, Sir3 has a winged-helix domain that can dimerize the protein. Sir4 has an N-terminal region of ∼750 amino acids that has DNA binding activity, but is not absolutely required for SIR function. Next downstream is the Sir2-Interaction Domain (SID), then a Partitioning and Anchoring domain (PAD) that makes contacts with nuclear periphery proteins, and the C-terminal coiled coil domain that dimerizes and makes interactions with Sir3.

A subset of molecular interactions that are important for SIR complex mechanism have been captured through high-resolution crystal structures; however, the nature by which these interactions cooperate within the full SIR complex to carry out its functions is not clear. The Sir3 BAH domain binds directly to the nucleosome via the histone H4 tail and multiple histones in the globular core of the nucleosome, captured by crystal structures[6–9]. The BAH domain is less than 25% of Sir3 and a small fraction of the full SIR complex, yet structures of the BAH-nucleosome offer the only major structural insights into SIR-nucleosome interactions. Structures of other domains of Sir3 and Sir2 and small fragments of Sir4 have also been solved[10–14]. Though these structures give less insight into the interactions with the nucleosome, they do provide reference points for biophysical approaches that can identify intermolecular interactions, such as cross-linking mass spectrometry[15].

Removal of histone modifications that promote transcription and open chromatin is essential for heterochromatin establishment in all organisms. In budding yeast, the heterochromatin histone code is the absence of nearly all modifications, while many organisms erase the same modifications as the SIR complex and add others, such as H3 lysine 9 and 27 methylation[16]. Sir2 is an NAD^+^-dependent lysine de-acetylase, which can remove many acetyl modifications to histone lysines[5, 17]. The SIR complex can bind directly to acetylated chromatin, engaging via Sir4-nucleosome interactions that are poorly-understood[5, 18]. Sir3 is bridged by Sir4 in this conformation, poised for action. Sir2 acts first, preferentially de-acetylating histone H4K16, which Sir3 then binds tightly to in the de-acetylated state. De-acetylation of additional histone lysines in the tails and core of the nucleosome by Sir2 facilitates chromatin compaction and reduces chromatin remodeling. Step-wise conformational changes during de-acetylation have been suggested by us and others[5, 19, 20], based on the two-step model for Sir3 interactions as it locks down onto the nucleosome after H4K16 de-acetylation. These changes may affect the orientation of Sir2 as it proceeds through erasure of all acetylation to facilitate Sir3-mediated compaction. At present, our models for these states of the SIR complex are coarse-grained and require improved methods for capturing these conformational switches.

We characterized the SIR heterotrimer by cross-linking mass spectrometry in solution. By doing so, we identified a novel protein-protein interface between Sir2 and the coiled-coil domain of Sir4. The coiled coil was previously demonstrated to associate with Sir3, suggesting this may be a hub of communication, connecting the early steps of the SIRc mechanism to the later. A key tyrosine in this interaction region of Sir2, conserved in nearly all eukaryotic sirtuins, is positioned to sense deacetylation state. In contrast, a pair of amino acids adjacent to the tyrosine may have co-evolved to interact specifically with Sir4 as a means of inter-subunit communication. By CRISPR genome editing, we mutated these two features of Sir2 and demonstrated that they each are critical for proper heterochromatin gene silencing. Our work highlights how a single multi-functional chromatin regulatory complex can stage a step-wise mechanism that requires a major transition in activities to achieve epigenetic gene repression.

## RESULTS

### Cross-linking mass spectrometry of the SIR complex reveals novel interactions

Because there is a lack of atomic-resolution structural information for the heterotrimeric SIR complex, we used cross-linking mass spectrometry to gain insight into specific protein-protein interactions, mapping inter-subunit cross-links between specific amino acids (Figure 2A). Purified SIR subunits with specific truncations to promote solubility and decrease higher-order interactions were used. Subunits were pre-incubated to form the complex and then subjected to cross-linking using the zero-length cross-linker DMTMM which cross-links lysines directly to aspartic and glutamic acid residues. Proteins were trypsinized and separated by strong cation exchange (SCX) chromatography to enrich for crosslinked peptides in specific fractions (Figure 2B) that were then subjected to tandem mass spectrometry (MS/MS). Cross-links were identified using pLink analysis software[21, 22].

**Figure 2.**
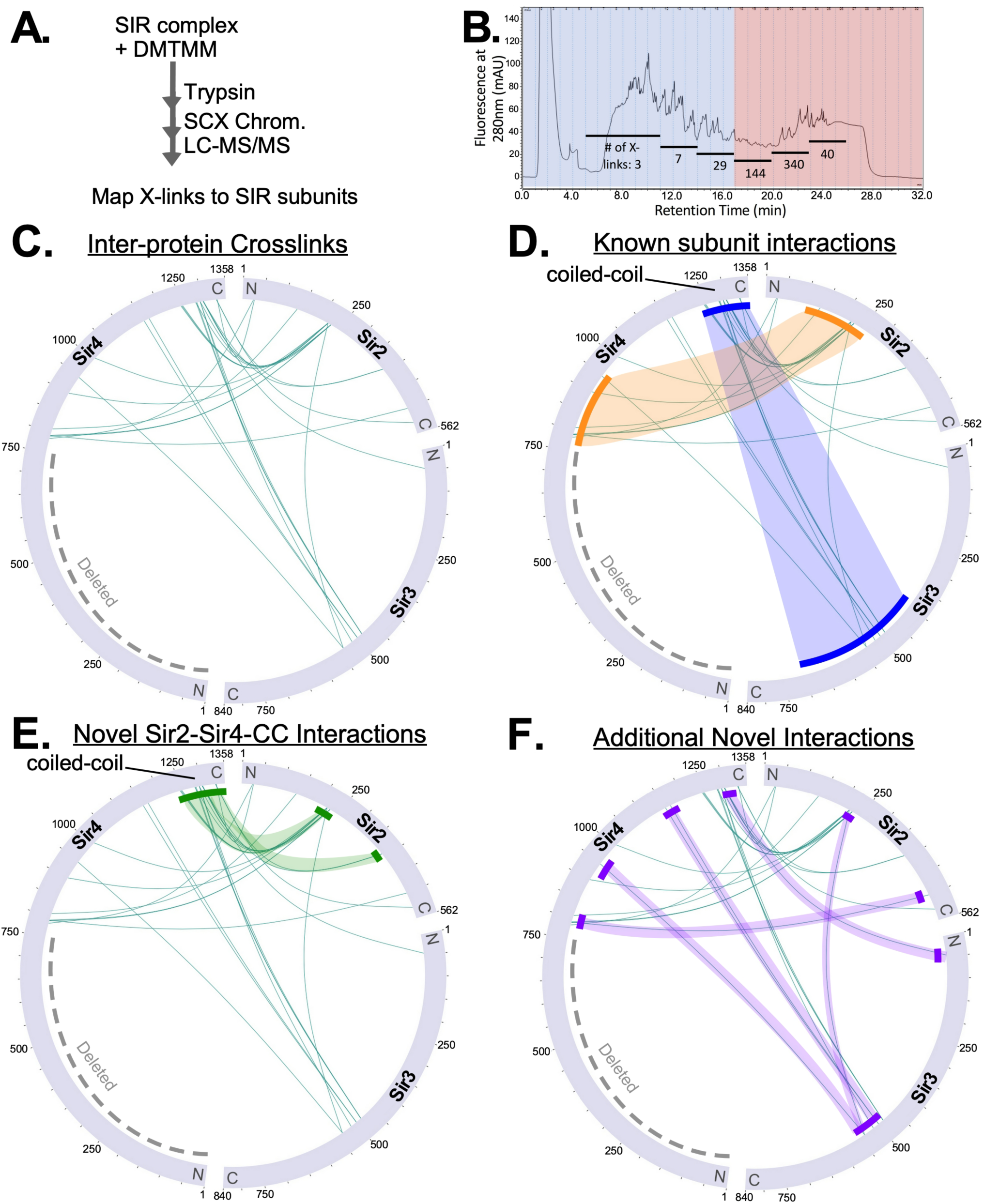
Crosslinking-mass spectrometry (XL-MS) identifies novel interactions between Sir2 and the coiled-coil of Sir4. A) Schematic for chemical crosslinking of the purified SIR complex using DMTMM, strong cation exchange (SCX), and tandem mass spectrometry (LC-MS/MS). B) Chromatogram of peptides eluted from the SCX column. The elution was split into six fractions which were individually analyzed by LC-MS/MS. Number of peptide crosslinks identified in each pool are indicated. C) Wheel diagram of all inter-protein crosslinks identified in the XL-MS analysis of the heterotrimeric SIR complex. Sir4-∆N and Sir3∆wh were used. D-F) The same wheel diagram as in C) with specific interactions highlighted: D) previously identified inter-subunit binding interfaces; E) novel interactions between Sir2 and Sir4; F) additional novel interactions identified, including a triangulation of position between Sir2, Sir3, and Sir4.

Mapping of peptide cross-links demonstrated many well-defined self cross-links within each protein, including identical peptide homo-meric cross-links of two identical peptides in the Sir4 coiled-coil (Sir4-CC), as expected for that dimerization region (Supplemental Figure S1). We focused our attention on the inter-protein cross-links to gain insight into the conformation of the SIR complex (Figure 2C). While the inter-subunit cross-links were more sparse than for intra-subunit, this is in keeping with the limited previously-identified interaction surfaces. We validated the known interactions between the Sir4 coiled-coil and the middle of Sir3(amino acids 485-515)[12] and also the interaction of Sir2 and the Sir2 Interaction Domain (SID) of Sir4 that promotes the Sir2/4 sub-complex[10] (Figure 2D). Unexpectedly, we also identified multiple peptide cross-links between Sir2 and the coiled-coil of Sir4 (Figure 2E), which have not been previously identified by genetic or biophysical approaches. The proximity of the novel Sir2 interaction on the Sir4 coiled-coil to the known Sir3 binding site suggests a potential hub of interactions within the SIR complex. This is supported by other novel cross-links between Sir2 and Sir3(Figure 2F), surprising because these two proteins have never been suggested to physically interact.

### Sir2 directly interacts with the coiled-coil domain of Sir4

We next determined whether the interaction we observed via crosslinking of Sir2 to the Sir4-CC was dependent on the known inter-subunit interactions within the SIR complex using two additional crosslinking mass spectrometry experiments. The proximity of residues on the Sir4-CC that crosslink to Sir2 and residues on the Sir4-CC that disrupts Sir3 interaction when mutated (Figure 3A), suggests either that Sir3 orients the Sir4-CC to interact with Sir2 or that the Sir4-CC is a potential hub of communication between activities of Sir2 and Sir3. We wished to test the first of these possibilities, whether the interaction with Sir3 may orient the Sir4 coiled-coil for interaction with Sir2, by omitting Sir3 in incubation of Sir4ΔN and Sir2 (Figure 3B,C). Second, we tested if the extensive interaction between the Sir4 SID and Sir2 was required for Sir2 to make additional contacts at the distal coiled-coil on Sir4, or whether the coiled-coil of Sir4 could bind Sir2 independently (Figure 3D). In both cases, we consistently observed cross-links between Sir2 and the coiled coil of Sir4, demonstrating a novel, independent interaction across this interface within the SIR complex.

**Figure 3.**
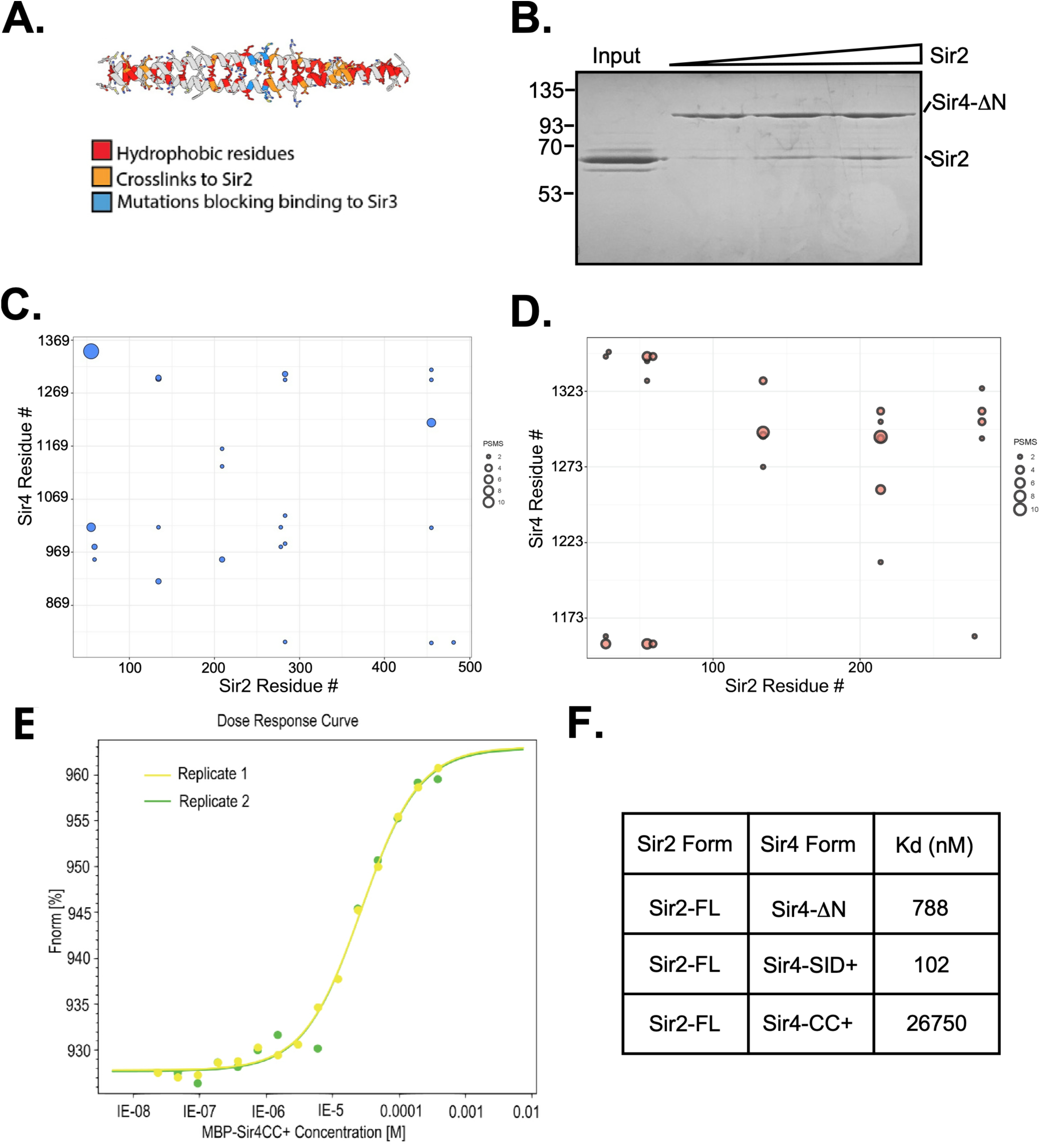
Sir2 interacts with the coiled coil of Sir4. A) Ribbon diagram of the dimer of Sir4 coiled-coil domain. Highlighted residues include those that were identified as Sir2-interacting by cross-linking mass spectrometry (orange) and those that, when mutated, disrupt Sir3 interaction (blue). B) Purified full-length Sir2 was incubated with MBP-tagged Sir4-∆N and the complex was isolated by amylose resin and resolved by SDS-PAGE and Coomassie stained. C) Matrix diagram depicting results from XL-MS of full-length Sir2 and Sir4-∆N, in the absence of Sir3. Sir2 axis is truncated due to no crosslinks occurring beyond the displayed sequence. Datapoint circle size is indicative of the number of peptide-spectrum matches (PSMs) at the given cross-link site. D) Matrix diagram depicting results from XL-MS of full-length Sir2 and Sir4-CC+. Sir2 sequence is truncated due to no crosslinks occurring beyond the displayed sequence. See methods for additional thresholding steps used for C) and D) analysis. E) Replicate binding curves from microscale thermophoresis (MST) experiments with fixed concentration of labeled Sir2 and MBP-Sir4CC+ titration range. F) Relative Kd measurements from MST experiments using the given constructs.

To characterize the interaction between Sir2 and the Sir4-CC further, we performed microscale thermophoresis (MST) experiments titrating the Sir4-CC into a solution of fluorescently-labeled, his-tagged Sir2 (Figure 3E). Binding between the two was observed, though the affinity was relatively low, compared to that of the Sir4-SID binding to Sir2 (Figure 3F). The lower affinity interaction of the Sir4-CC with Sir2 suggests that this interaction may not be a primary driver of association of the two full-length proteins, but instead may occur intermittently, perhaps in a regulated manner.

### Identification of Sir2 279-286 as a novel region that is important for silencing *in vivo*

We next aimed to determine whether the interaction between Sir2 and the Sir4-CC may be important for the regulation of gene silencing in *S. cerevisiae*. We used a previously-designed yeast strain bearing GFP and RFP reporter genes at the *HMR* and *HML* loci, respectively[23, 24] (Figure 4A). When the SIR complex activity is disrupted, both reporters are de-repressed and the colonies and cells fluoresce. CRISPR genome editing was used to make markerless edits to the *SIR* genes to mutate residues within major cross-linking regions, assessed by high frequency of cross-links and thresholding.

**Figure 4.**
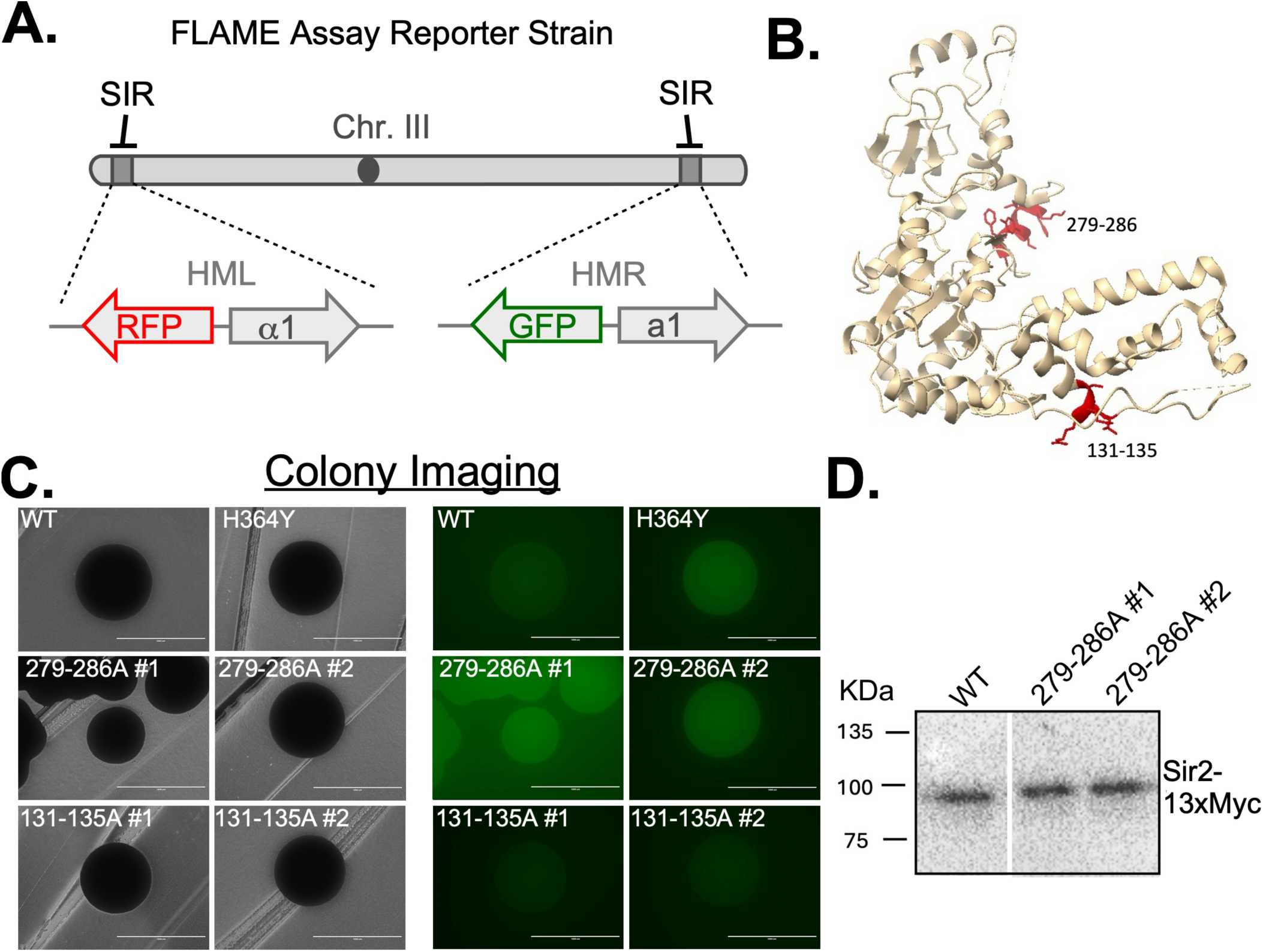
Identification of a novel regulatory region of Sir2. A) Schematic describing the FLAME assay system, previously developed [23, 24]. RFP and GFP are each substituted B) Ribbon diagram representation of Sir2 (PDB 4IAO) structure with two highlighted patches (red) that both make multiple crosslinks to the Sir4 coiled coil. C) Image of yeast colonies on the transillumination and GFP channels using the FLAME system strain. WT and Sir2 catalytic mutant (H364Y) are used as reference to image multiple isolates of strains mutated to alanine via CRISPR/Cas9 at Sir2131-135 and Sir2 279-286. One millimeter scale bar shown. D) Western analysis of myc-tagged Sir2 strains comparing wild-type and two Sir2 279-286A mutant isolates.

Two regions of Sir2 were the primary focuses of the mutational analysis, residues 131-135 and 279-286 (Figure 4B), neither of which had been previously identified as interacting with another protein and no mutations with phenotypes had been previously reported for either region. We first created mutant strains in which all amino acids in either region were mutated to alanines. The mutation of amino acids 279-286 led to de-repression of the HM loci by colony fluorescence, comparable to a mutation of Sir2 that abolishes the deacetylation activity, H364Y [25] (Figure 4C). We also confirmed that this de-repression was not caused by overall destabilization and degradation of the protein (Figure 4D). Conversely, mutating 131-135, nor in other crosslinking sites, did not lead to significant de-repression and fluorescence (Figure 4C, Supplemental Figure S2).

### Sir4-CC interacts with a conserved sensor region of Sir2

The Sir2 residues that interact with the Sir4-CC are adjacent to the highly-conserved cofactor binding loop (CBL) of Sir2, widely found in sirtuins [26, 27] (Figure 5A,B). The CBL adopts many conformations in the available sirtuin crystal structures, typically depending on the state of cofactor binding. NAD+-bound sirtuin structures tend to have a closely-packed, ordered CBL that makes explicit contact with the cofactor bound, while crystal structures without cofactor bound typically have less ordered or flexible and unresolved CBLs (Figure 5C). The adjacent region to the CBL, the stretch of amino acids that includes the specific Sir4-CC cross-link site at Sir2-K283, which we henceforth term the cofactor binding loop extension, is itself highly-conserved in a subset of sirtuins, from yeast to humans (Figure 5A). For example, human sirtuins SIRT1, 2, 3, 4, and 5 all have a conserved tyrosine, corresponding to Y281 in *S. cerevisiae*, yet SIRT6 and SIRT7 do not. This tyrosine is conserved in *S. cerevisiae* sirtuins Hst1,2, and 3, but not Hst4. The CBL extension has conserved residues that, in some sirtuin crystal structures, point towards and/or make hydrogen bonding interactions with the ribose or nicotinamide moiety of NAD+. Y281 (in ScSir2) in particular has been observed to do this, for example in the Sir2/Sir4-SID crystal structure with ADP-ribose bound [10], and a structure of ScSir2 in the act of deacetylation also has Y281 pointing toward the deacetylation intermediate. The conservation of certain residues within the CBL extension and the physical interaction that it makes with Sir4CC is highly suggestive of a means to couple the sensing of NAD+, by residues such as Y281, with communication to the Sir4CC, and possibly to its other binding partner Sir3.

**Figure 5.**
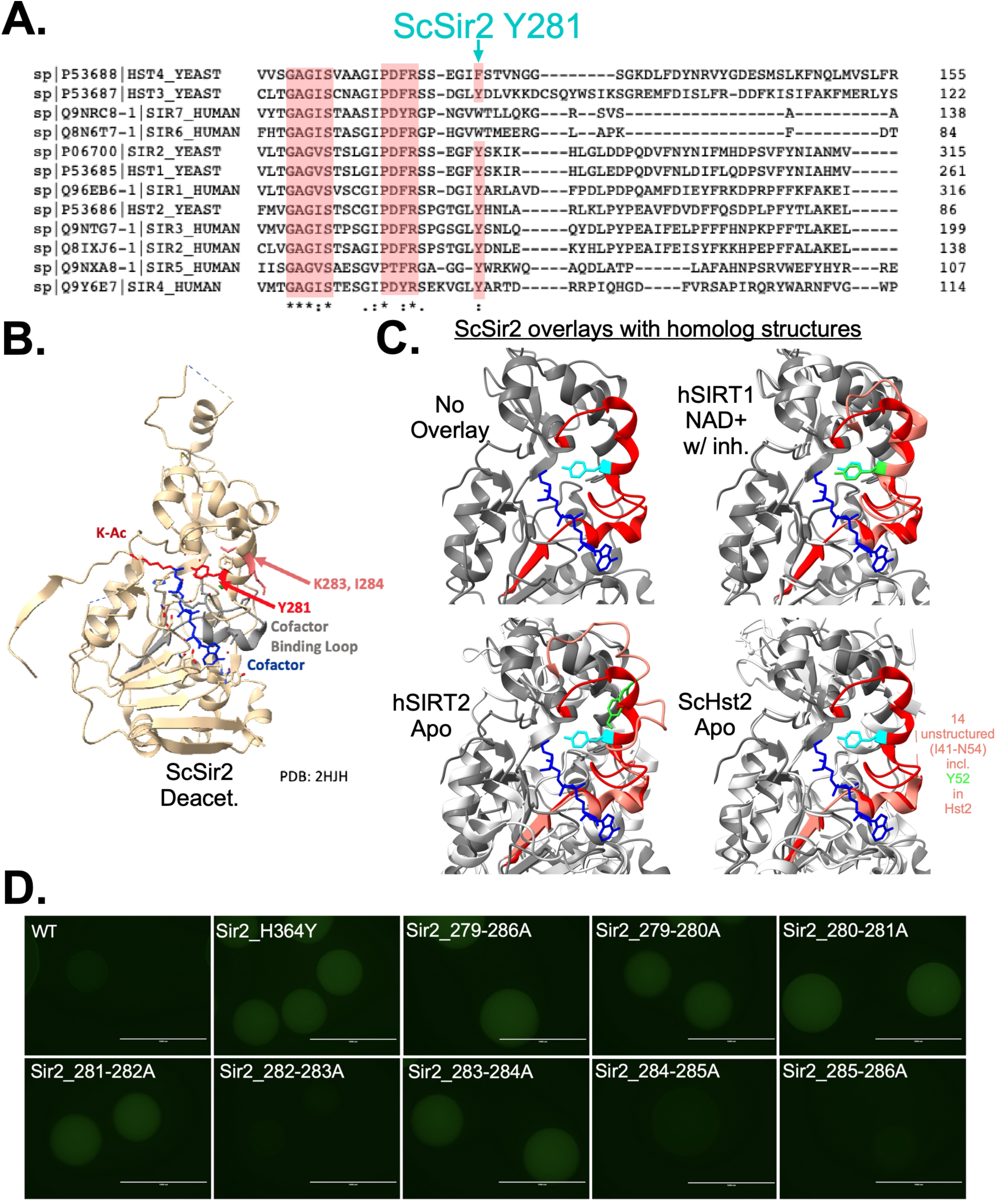
Conservation and analysis of key residues in the Sir2 cofactor binding loop extension. A) Alignment of all yeast and human sirtuins, highlighting the conservation in the cofactor binding loop extension and in the tyrosine residue at position 281 in budding yeast Sir2. B) Structure of budding yeast Sir2 (ScSir2) in the act of deacetylation with cofactor intermediate (blue) covalently attached to the lysine being deacetylated (K-Ac). Key residues important for ScSir2 silencing (Y281, K283, I284) highlighted in red. Unpublished PDB deposition 2HJH. C) Overlay of the 2HJH structure and structures of other sirtuins in the apo and nucleotide bound forms, demonstrating the dynamics of the cofactor binding loop (dark red for 2HJH, light red for other sirtuin) and the tyrosine sensor (ScSir2 Y281 in cyan, other sirtuin tyrosine in green). Structures with a variety of cofactor binding status were used. The cofactor binding loop extension of ScHst2 (*lower right*) is too flexible and is not resolved in the structure. PDB references: 4i5i Chain A (hSIRT1), 1j8f Chain A (hSIRT2), 1q14 Chain A (ScHst2). D) Combinations of double mutants within Sir2 279-286 were assessed by FLAME GFP assay. One millimeter scale bar shown.

### Mutational dissection reveals two essential components for silencing in the Sir2 279-286 region

To determine which residues are important for silencing within the 279-286 region, we began by mutating each tandem two amino acid pair and assessing colony fluorescence. We observed specific pairs with increased fluorescence, centering on Sir2 residues G279, F280, Y281, as well as K283 and I284 (Figure 5D).

While the colony-based fluorescence was useful for screening purposes, we next turned to a more quantitative readout, using single-cell imaging. This approach allows us to not only provide a more accurate fluorescence intensity quantification but also profiles many cells in the same genetic background, to determine if there is more than one distinct population. For example, variegation in the population can lead to a bimodal distribution of cells, some of which have repressed GFP while other cells have de-repressed HM loci and subsequently the fluorescent reporters. Using this single-cell method, we were able to quantitatively distinguish strains that repressed the HM loci versus those where SIR-based repression was disrupted. We first characterized the mutant in the 279-286 patch, which exhibited de-repression of the HM loci, as expected, by visual inspection of single cells by fluorescence microscopy (Figure 6A). Quantification demonstrated that Sir2 279-286A cells have fluorescent signal similar to the catalytic mutant Sir2 H364Y, slightly lower than a full deletion of Sir2. Single-cell quantitative profiling of the double mutants in the Sir2 279-286 patch demonstrated that Sir2 280-281A and 281-282A mutants were de-repressed at HM to the same extent as the Sir2 deletion mutant (Figure 6B). Since both of these double-mutants contained the Sir2 Y281A mutation, we profiled this single mutant and found that it displayed the same phenotype as the Sir2 delete. A similar strong phenotype was observed for the Sir2 283-284A double-mutant, although other double-mutants containing one, but not the other, of these two mutations did not produce a phenotype, suggesting that these two residues are together important for SIR-dependent repression.

**Figure 6.**
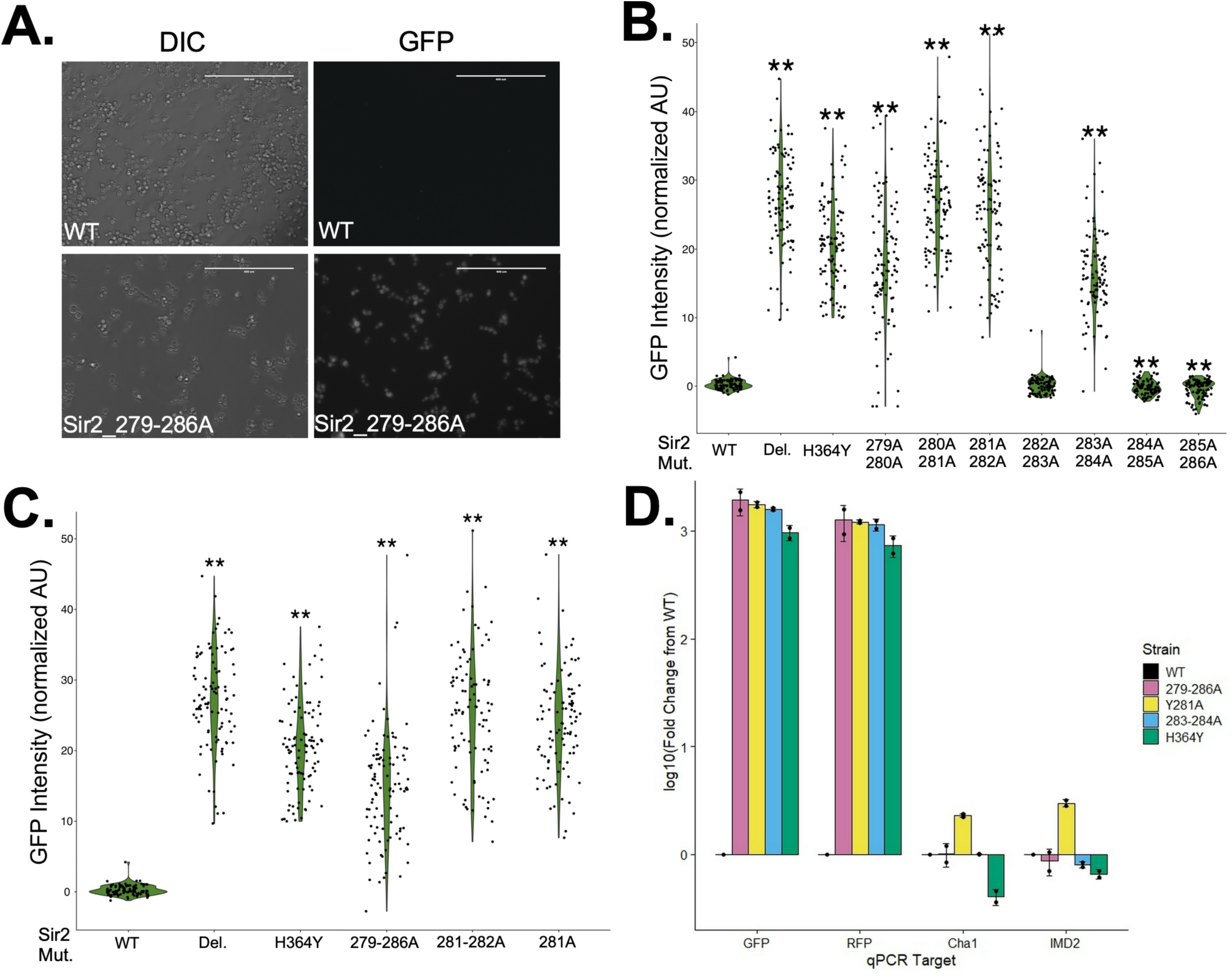
Sir2 Y281 alone and K283/I284 together are critical for SIR-dependent gene repression. A) Example of microscopy images, contrast and GFP channel, depicting WT and Sir2_279-286A FLAME strains. B) Single-cell quantification of GFP intensity for double mutants in the 279-286 patch of Sir2. C) Analysis of Sir2 Y281. ** over plots indicate t-test against WT yielded a p-value below 0.01. D) RT-qPCR determination of mRNA changes for SIR-regulated loci in indicated Sir2 mutants, log10 fold-change over WT from biological replicate strains. GFP and RFP genes located within HMR and HML, respectively.

As an orthogonal quantitative technique to determine the effect of Sir2 mutations, we use RT-qPCR for the GFP and RFP reporter genes and in addition assayed for expression of two subtelomeric genes that were previously reported to be de-repressed when *SIR* genes were deleted [28]. Compared to WT, the Y281A and 283-284A mutants both displayed a loss of silencing, resulting in >1000-fold upregulation of the GFP and RFP reporter genes in the *HM* loci, comparable to the catalytic mutant of Sir2 (Figure 6D). Interestingly, the subtelomeric genes *CHA1* and *IMD2* were only de-repressed in the Sir2 Y281A mutant.

## DISCUSSION

We set out to more fully map the protein-protein interactions that occur within the SIR complex and identified a novel interaction interface that is required for SIR heterochromatin, potentially via coordinating a transition from the early steps in the mechanism to the later steps. Using cross-linking mass spectrometry of the SIR complex allowed us to identify these novel interactions between Sir2 amino acids 279-286 and the Sir4 coiled-coil domain, which were further supported by microscale thermophoresis of Sir2 with Sir4 coiled-coil. Subsequently, we mutated these regions by CRISPR editing of the endogenous *SIR* genes to uncover the Sir2 279-286 region as an important regulator of SIR-dependent epigenetic gene repression. These novel connections between Sir2 and the region on Sir4 that is already known to bind Sir3 suggest a mode of coordination between the early events of Sir2 deacetylation of nucleosomes and the subsequent stable binding of those deacetylated nucleosomes by Sir3 that converts the chromatin to a compact, repressed state. Our work helps uncover how a single multi-functional heterochromatin complex, sufficient to carry out all the core steps of epigenetic gene repression, can coordinate diverse activities.

### Triangulation of the SIR complex subunits by the Sir4 coiled-coil

Using cross-linking mass spectrometry of the SIR complex allowed us to identify a previously-uncharacterized interaction between Sir2 and the Sir4 coiled-coil. This interaction is not critical to Sir2-Sir4 association, due to the higher-affinity between Sir2 and the central Sir2-interacting domain (SID) of Sir4, which is stable enough to be co-purified and co-crystallized [10]. The nature of this weaker Sir2-Sir4 coiled coil interaction suggests that it may be dynamic, offering the opportunity for conformational change within the SIR complex where Sir2 can bind to the Sir4 coiled-coil transiently, potentially at a specific step in the mechanism of heterochromatin formation. The proximity of the regions of the Sir4 coiled-coil that interact with Sir2 and those that interact with Sir3 suggest that a dynamic mode of Sir2 binding could be transmitted to Sir3 through the Sir4 coiled-coil. In support of the potential for coordination, we identified an additional novel Sir2-Sir3 interaction (Figure 2F), the first to be discovered between these two subunits. While this singular point of crosslinking between Sir2 and Sir3 may not be evidence of a significant subunit interface that can directly influence the SIR mechanism, it provides further evidence in favor of the Sir4 coiled-coil as a triangulation hub between the subunits, perhaps as a conduit of information between Sir2’s dynamic deacetylation mode and Sir3’s stable lockdown mode. Finally, it is notable that the Sir2 279-286 region that is critical for gene repression was crosslinked to the Sir4 coiled-coil only in the absence of Sir3, suggesting that there may be some level of influence that can shift the interactions.

### Coordination of the Sir2 cofactor binding loop extension residues as a potential mode-switching mechanism for the SIR complex

To our knowledge, no mutant from S277-L286 has been isolated in a screen or characterized individually for silencing activity or Sir2 activity. Mutants of F274, R275, and S276, the last residues of the CBL, have been characterized as silencing mutants and demonstrate a variety of effects on Sir2 activities [29–31] and CBL deletions lead to loss of silencing [32]. R275 is deficient for silencing, but in vitro histone peptide deacetylation is not affected [29]. Fusions that replace the catalytic components of Sir2 with fairly conserved regions from other species frequently fail to complement in gene silencing [32, 33]. While the CBL extension has some conserved residues, others, such as the critical K283 and I284 combination, are not conserved, suggesting that the coupling to Sir4 is lost by these fusions. Sherman et al. showed that the swap of the human SIRT2 catalytic domain into yeast Sir2 is dominant negative, suggesting that the mutant could bind to Sir4 but not carry out the full mechanism; rDNA silencing was intact in the fusion [32]. Mead et al. demonstrated that swapping singular regions of Sir2 into Hst1 was not sufficient to restore SIR silencing; however no combinations that included the main Sir4 binding domain and the region around the CBL were constructed. The regulatory mechanism involving the extended cofactor binding loop in Sir2 depends on NAD+, since a catalytic domain that does not depend on NAD+ subverts this regulatory mechanism [34]. These previous findings support the importance of the novel Sir2-Sir4CC interaction that we have identified in driving NAD+-dependent formation of SIR heterochromatin.

With our current study and in light of the previous work outlined above, we propose a model in which Sir2 Y281 senses active site occupancy and this is communicated via residues K283 and I284 that make direct interactions with the Sir4 coiled-coil, preventing the “lockdown” stage of the mechanism (Figure 7). Once deacetylation is complete, the sensor is required to switch to the next mode. This may explain why a Y281A sensor mutant displays a stronger phenotype, de-repressing subtelomeric genes, by preventing the licensing step. The 283-284A mutant may simply bypass the sensor, removing the hold on the “lockdown” stage, allowing from some repression to occur. The wild-type function of this staging could be important for the tight, >1000-fold repression at the *HM* loci, but dispensable for tuning subtelomeric genes. When both elements are mutated in the 279-286A mutant, the 283-284A mutation suppresses the Y281A mutation by allowing the “lockdown” stage to occur, even in the absence of the sensor that stages the mechanism for the strongest repression.

**Figure 7.**
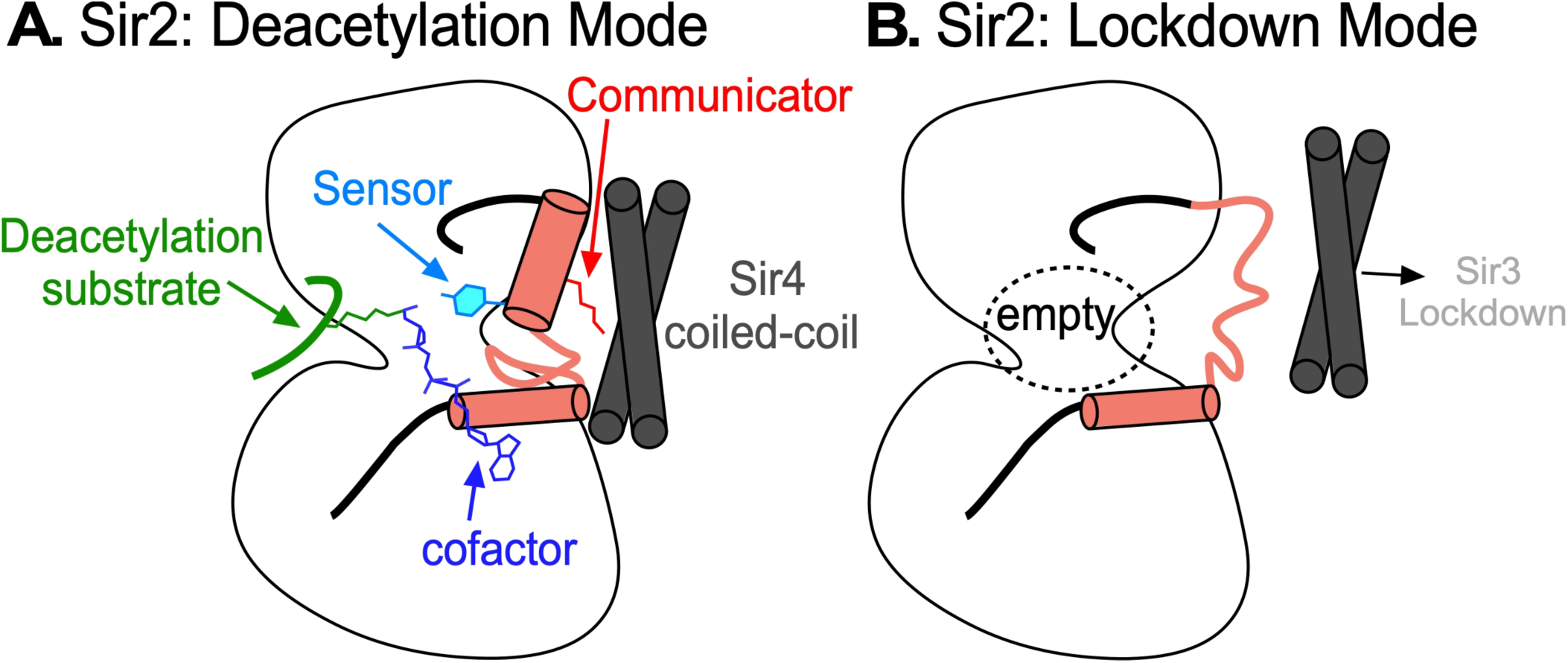
Proposed model for Sir2 conformational changes driving the two modes of the SIRc mechanism via communication with the Sir4 coiled-coil. A) In Deacetylation Mode, Sir2 is engaged with an acetylated histone substrate, using the NAD+ cofactor (blue) to drive removal of acetyl marks on histone lysine residues. In this state, the sensor, ScSir2 Y281 (cyan), detects the activity in the catalytic site, which promotes ordering of the helix that Y281 is on, the cofactor binding loop extension (light red), and orients the communicator residue(s), Sir2 K283 (red), which directly contacts the Sir4 coiled-coil to promote this mode of action. B) Once deacetylation is compete, the Sir2 catalytic site is empty, disengaging the sensor, which then leads to a disordered CBL extension, communicating to Sir4 that Sir3 can lockdown onto chromatin.

## Conclusion

Our work using cross-linking mass spectrometry as a means of probing the SIR heterotrimer led to identification of inter-subunit physical interactions that had not be previously identified, despite a rich history of SIR genetics and biochemistry. The specific Sir2-Sir4 interface we characterize sits poised to sense completion of deacetylation and license the second half of them mechanism where Sir3 locks down on chromatin to repress transcription. This mechanism may be conserved as a means to stage deacetylation that is coupled to secondary coordination within many sirtuin complexes with diverse functions.

## METHODS

### Sir Protein Purifications

Recombinant Sir3 was purified as described in [5], via FLAG purification for both full-length and winged helix deletion (Sir3 1-850). Sir2 (untagged final) was expressed in E. coli as a GST fusion and purified by glutathione affinity chromatography and PreScission Protease cleavage. Sir2-6XHis was expressed in E. coli and purified by via Ni-NTA and MonoQ chromatography steps and finally dialyzed into 30 mM Hepes pH 7.5, 100 mM NaCl, 5% glycerol. Sir4 constructs were all cloned downstream of maltose-binding protein (MBP) and expressed in E. coli. MBP fusions were purified via amylose resin chromatography and elution with maltose. Sir4(744-1358, “Sir4-∆N) and Sir4(744-115, Sir4-SID+) were further purified by MonoS chromatography. All Sir4 preps were dialyzed into 50 mM Hepes pH 7.5, 100 mM NaCl, 1 mM DTT, 5-10% glycerol.

### Crosslinking Mass Spectrometry

Crosslinking was performed at neutral pH, 35°C for 1hr and quenched with Ammonium bicarbonate. Trypsin peptides were separated by strong cation exchange chromatography where the crosslinked peptides with higher total charge generally elute later from the gradient (Fig. 2B). Tandem mass spectrometry (MS/MS) analysis was run and cross-links were identified using a data-dependent acquisition method on a Fusion-Lumos. The method was optimized and biased towards peptides found in higher charge states(z > 2+). Cross-links were further identified using pLink analysis software[21, 22]. Analysis of the SIR complex used E. coli-derived Sir2 and a large fragment of Sir4(744-1358) as a maltose binding protein fusion. To minimize oligomerization for XL-MS, to better define unique protein-protein interactions, a Sir3 construct with deleted winged helix dimerization domain was used, purified as described yeast overexpression [5]. This left the Sir4 dimeric coiled-coil region (Sir4cc) at the C-terminus as the single self-interaction domain, which cannot be deleted without disrupting Sir3 interactions in the SIR complex[35]. Circular crosslink figures were generated with xiView (https://xiview.org/) with a minimum match score of 27. Experiments in Figure 3 were further thresholded by masking any Sir2 or Sir4 site that was also frequently crosslinked non-specifically to maltose binding protein in the Sir4 fusion constructs (Supplemental Table S1). Experiments in Figure 3 were displayed as a matrix for added resolution. All data are publicly available on the Proxl server; http://www.yeastrc.org/proxl_public/projectReadProcessCode.do?code=hmj2tvem8kxpnrn0vt8o995yx8aeltd

### Microscale thermophoresis (MST)

MST was performed on a NanoTemper Technologies Monolith NT.115 pico instrument, medium power, auto-detect excitation, Pico-RED excitation color with RED-tris-NTA labeling of Sir2-6Xhis, 24-25°C, for all experiments. All experiments performed with constant Sir2-6Xhis in 30 mM Hepes pH 7.9, 100 mM NaCl. 10 nM Sir2-6Xhis was used for Sir4(744-1358, “Sir4-∆N) and Sir4(744-115, Sir4-SID+) and 50 nM for Sir4(1150-1358, Sir4-CC+). MO Affinity Analysis software was used for binding curve fitting and Kd value determination.

### Amylose Bead Pulldown

Untagged Sir2 was incubated with MBP-tagged Sir4 constructs at 16°C for 1 hr in Binding Buffer (50 mM Hepes pH 7.9, 200 mM NaCl, 10% glycerol, 1 mM DTT, 0.1% Tween-20). 50 µL amylose resin was added and incubated at 16°C for 1 hr on a Thermomixer at 1000 rpm. Beads were collected at 4°C by 1 min centrifugation at 500 x g, washed with 75 µL Binding Buffer, then protein was eluted in the wash buffer with 10 mM maltose by 5 min. mixing at 23°C and 1 min centrifugation at 500 x g 4°C. Samples were mixed with 2X Bio-Rad SDS Sample Buffer and run on a 12% polyacrylamide gel with Coomassie stain.

### CRISPR genome editing

Plasmid pML104 (Addgene #67638) was used for CRISPR-Cas9 yeast genome editing as previously described [36]. pML104, containing Cas9 and an sgRNA cassette, was digested with SwaI and BclI, then ligated with annealed oligonucleotides to serve as the guide sequence for the sgRNA. Donor DNA was with the target mutation, along with a silent PAM site mutation to prevent re-cutting of the genomic DNA, was produced by PCR from previously-generated plasmid or by gBlock synthesis (IDT). Donor DNAs included ∼40-70 bp of homology on either side to promote homologous recombination DNA repair. Where necessary, silent mutations between the Cas9 cut site and the target mutation were made to limit recombination before the target editing site. Yeast strains were transformed with 1 μg of pML104 and 3 μg of donor DNA using lithium acetate and PEG. Mutants were confirmed by extracting genomic DNA with lithium acetate and SDS, amplifying the target gene, and sequencing. Cells were counter-selected on 5-fluoroorotic acid (5-FOA) plates to remove the pML104 plasmid. All yeast strains generated listed in Supplemental Table S2.

### FLAME assays by colony

Yeast colonies were grown from glycerol stock on synthetic complete medium plates for 2-3 days. Colonies were imaged on an AMG EVOS imager in transillumination and GFP channel using 30% fluorescence threshold at 4X magnification.

### Single cell FLAME assays

Yeast strains were grown in synthetic complete medium with glucose to 0.3-0.5 OD, transferred to 96-well plates and imaged on an EVOS microscope at 40X objective for transmission, GFP, and RFP collection. Images were analyzed by FIJI. First, transmission images were used to make a mask of >100 individual cell outlines, along with background outlines. The mask is then used to collect intensity data for GFP and RFP images. Background-subtracted values were visualized in violin plot format with Student’s t-test for statistical significance.

### RT-qPCR

RNA was extracted from yeast using hot acid phenol and SDS, followed by a wash with chloroform and ethanol precipitation. Samples were treated with Turbo DNase and cleaned up with the Qiagen RNeasy Mini Kit. cDNA was produced using the Applied Biosystems High Capacity cDNA Reverse Transcription Kit. qPCR was performed using the SYBR GreenER qPCR SuperMix Universal (Thermo) under standard conditions using the primers in Supplemental Table 3.

## Acknowledgements

This work was funded by the NSF (Research Grant 2002231 from MCB to A.M.J and Graduate Research Fellowship 25A9567 to E.J.M.), the NIH (R35GM144358 to A.M.J.), and NCI Cancer Center Support Grant P30CA046934 to the University of Colorado Cancer Center. We would like to acknowledge Jacki Williams, Cassie Smith, Ari Rivera, Nahili Mohammed, Kim Miranda, and Aaron Issaian for technical assistance, Jasper Rine and his lab and John Wyrick and his lab for generous gifts of strains.

## Conflict of Interest

The authors declare that they have no conflicts of interests with the contents of this article.

## Author Contributions

J.K., E.J.M. and A.M.J. designed the research project; J.K., E.J.M., M.N., N.S., L.S., J.S., and N.H generated reagents; J.K., E.J.M., M.N., N.S., and L.S. performed experiments and analyzed preliminary data. J.K., E.J.M., K. H., and A.M.J. synthesized the data and wrote the manuscript.

**Supplemental Table S1.** Crosslinked amino acids identified in experiments with Sir2 and constructs of Sir4, identified with high-stringency filter.

**Supplemental Table S2.** Yeast strains used in this study.

**Supplemental Table S3.** RT-qPCR primers used in this study.

---

**Supplemental Figure S1.**
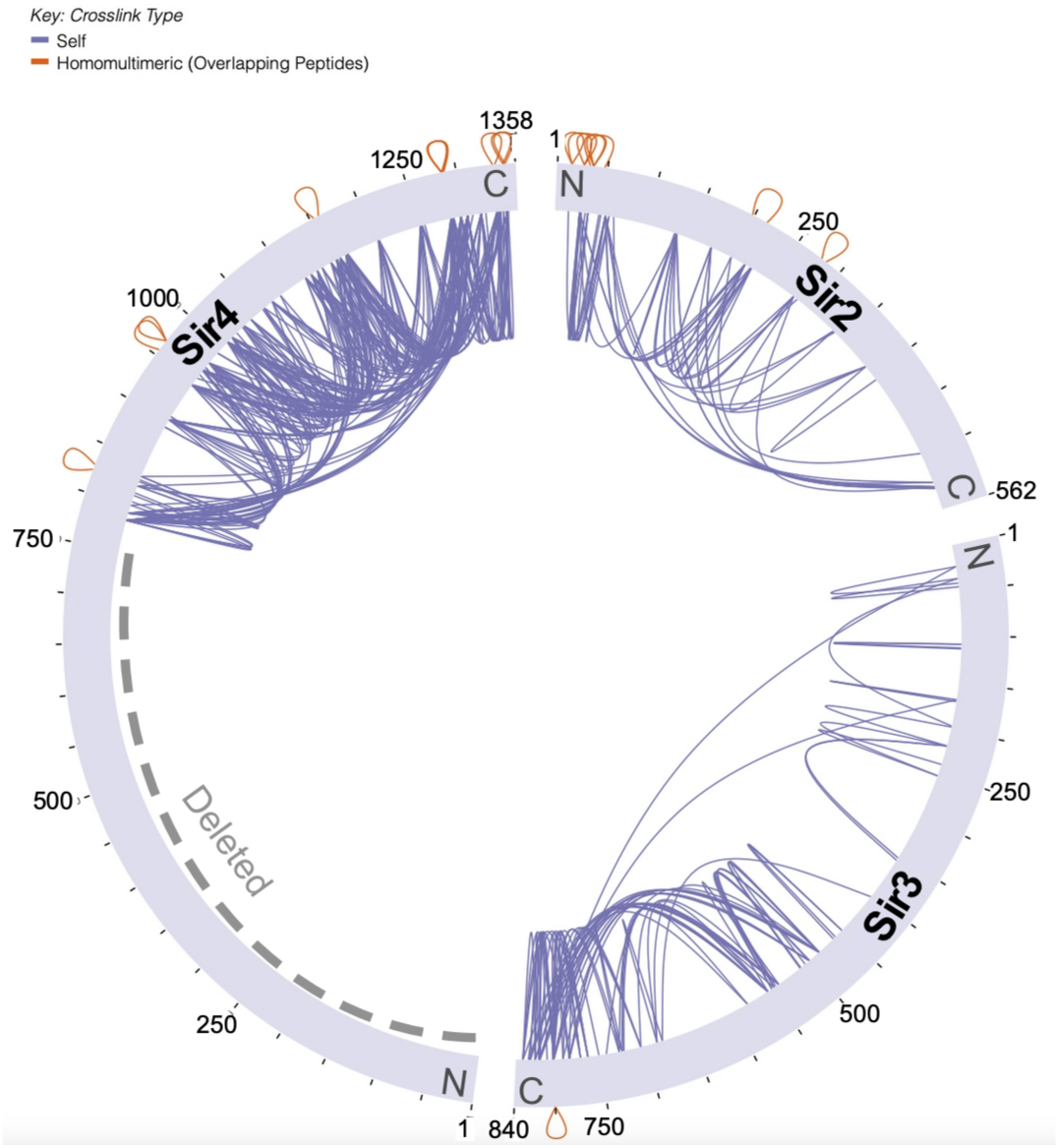
Self crosslinks within the SIR complex. A) Example SIR complex assembly with Sir4∆N, Sir3, and Sir2 incubation and elution with maltose. B) xiView schematic depicting the crosslinks within each Sir protein construct, including crosslinks with different parts of the protein (Self, purple) and two crosslinks between identical peptides (Homomultimeric, orange).

**Supplemental Figure S2.**
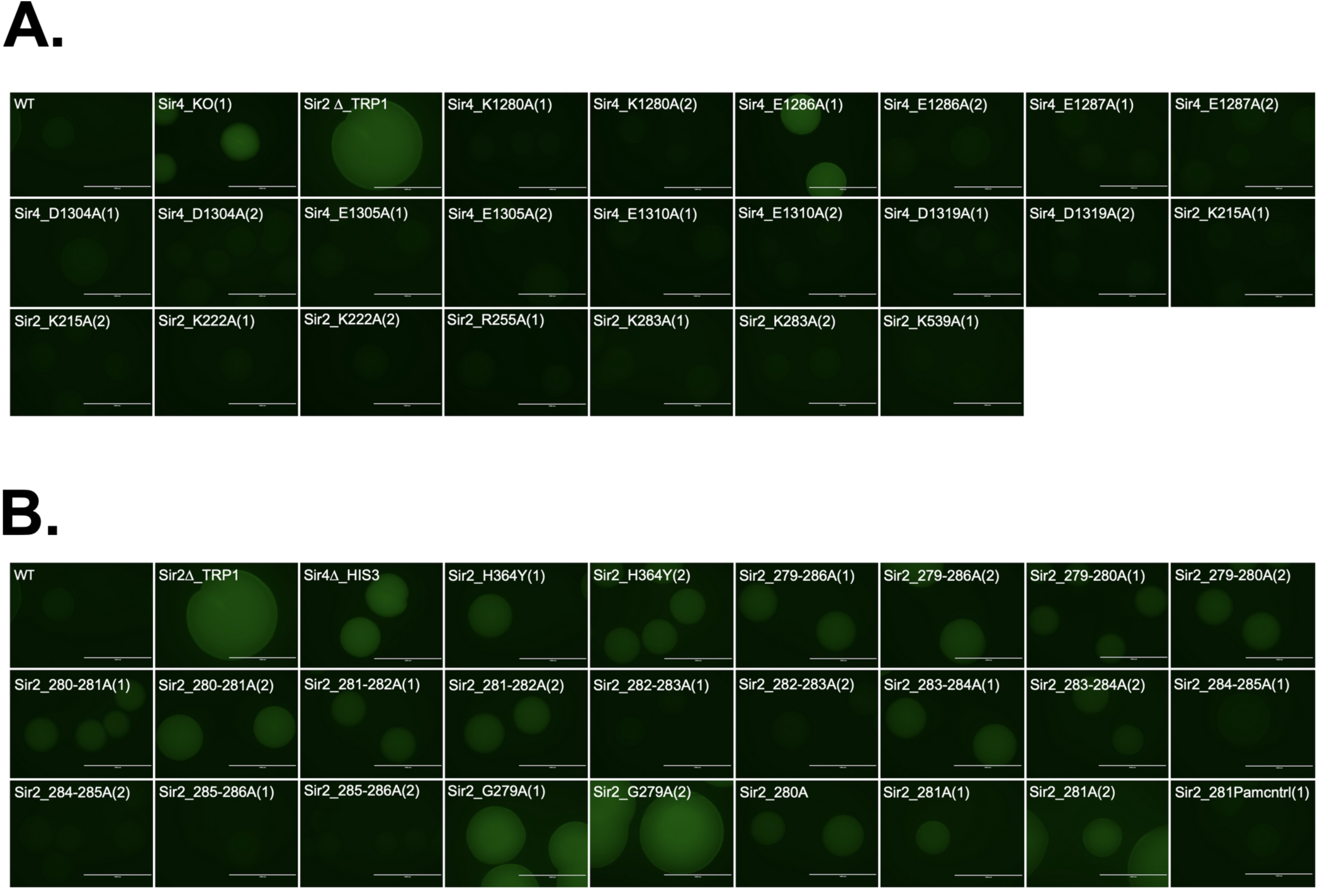
Additional Colony FLAME Assay Results. A) GFP channel images from FLAME strains with alanine mutations of crosslink sites in Sir2 and Sir4. B) GFP channel images of FLAME strains of Sir2 mutations, including multiple isolates and a control for silent mutations of the Sir2 279-286 regions that includes the PAM site mutation used for all strains in the figure panel.

**Supplemental Figure S3.**
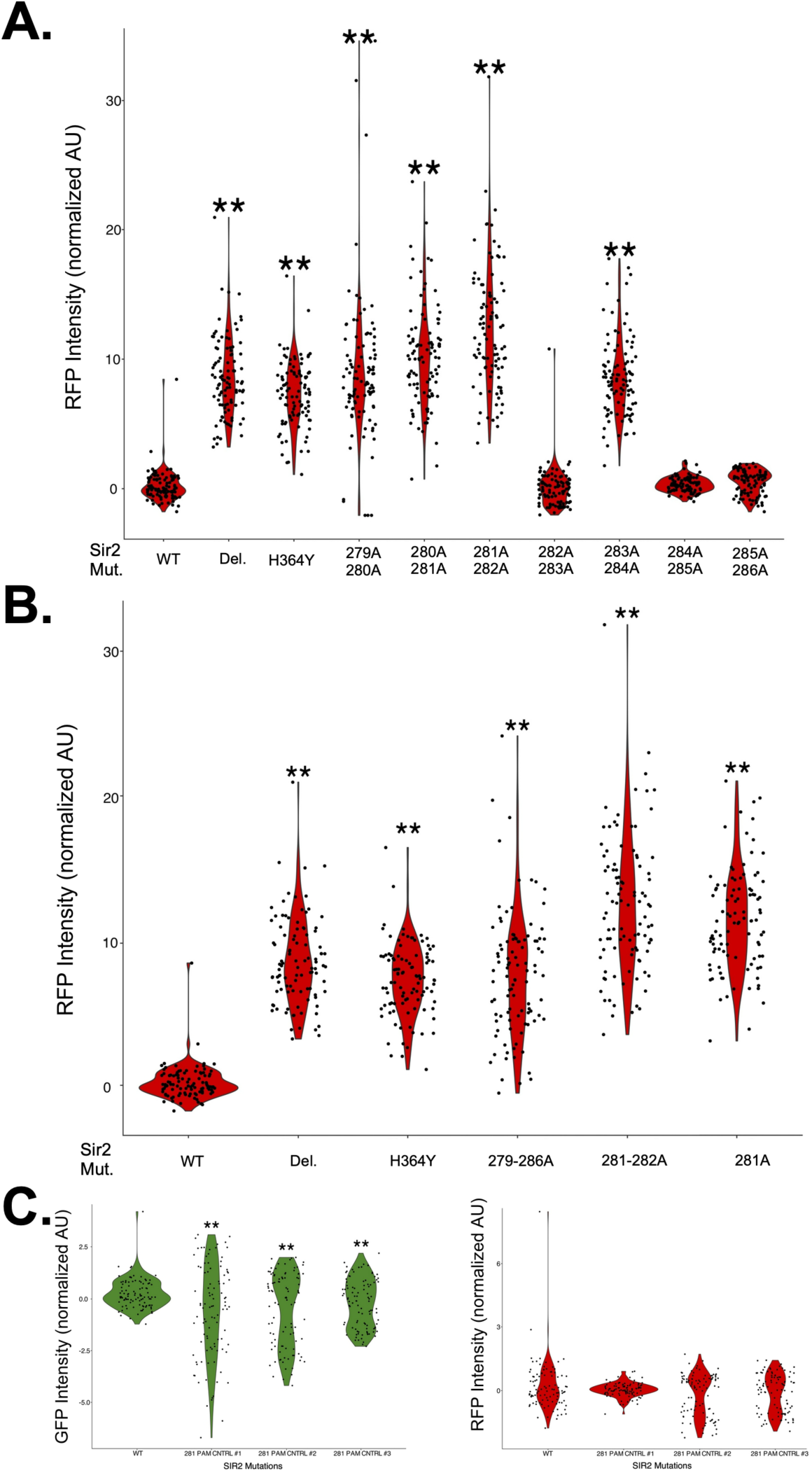
Additional Single-Cell FLAME Assay results. A,B) RFP channel single cell FLAME results for indicated Sir2 mutants. C) GFP and RFP channel results for three strains with CRISPR-engineered PAM mutation only at the site used for mutation of Y281.

## Notes

### Competing Interest Statement

The authors have declared no competing interest.

### Summary of Updates

Text modified for clarity, Figure 6D and Supplemental figures added.

